# Quantitative, model-based estimates of variability in the serial interval of *Plasmodium falciparum* malaria

**DOI:** 10.1101/058859

**Authors:** John H. Huber, Geoffrey L. Johnston, Bryan Greenhouse, David L. Smith, T. Alex Perkins

## Abstract

**Background**: The serial interval is a fundamentally important quantity in infectious disease epidemiology that has numerous applications to inferring patterns of transmission from case data. Many of these applications are apropos to efforts to eliminate *Plasmodium falciparum (Pf)* malaria from locations throughout the world, yet the serial interval for this disease is poorly understood quantitatively.

**Results**: To obtain a quantitative estimate of the serial interval for *Pf* malaria, we took the sum of components of the *Pf* malaria transmission cycle based on a combination of mathematical models and empirical data. During this process, we identified a number of factors that account for substantial variability in the serial interval across different contexts. Treatment with antimalarial drugs roughly halves the serial interval, seasonality results in different serial intervals at different points in the transmission season, and variability in within-host dynamics results in many individuals whose serial intervals do not follow average behavior.

**Conclusions**: These results have important implications for epidemiological applications that rely on quantitative estimates of the serial interval of *Pf* malaria and other diseases characterized by prolonged infections and complex ecological drivers.

## BACKGROUND

The basic reproduction number *R*_0_, defined as the expected number of secondary cases arising from a single primary case in a susceptible population, is well known and of fundamental importance in infectious disease epidemiology. Despite extensive efforts to model, measure, and map *R*_0_ globally for *Plasmodium falciparum* (*Pf*) malaria [1, 2], little has been done to quantify its temporal analogue, the serial interval. Defined as the time between the clinical presentation of primary and secondary cases, the serial interval is also of fundamental importance [3]. Probabilistic descriptions of the serial interval provide a basis for identifying sources of infection [4], for assessing whether cases are causally linked [5, 6], for analyzing incidence data to estimate temporal variation in transmission and its environmental drivers [7, 8], and for determining whether a pathogen can be declared eliminated [9].

For directly transmitted diseases, the serial interval can be measured through contact tracing or with household data [10, 11]. For malaria and other mosquito-borne diseases, this would require the impossible task of tracing mosquito blood meals between people, so the serial interval must be estimated indirectly. Case data have been analyzed with scan statistics to estimate the serial interval (e.g., [12]), but the resolution of these estimates is extremely crude. Scan statistics are generally not capable of capturing heterogeneity in the serial interval distribution across different contexts, but heterogeneity in the ecology of *Pf* malaria across space, time, urban-rural gradients, and other respects is an important feature of its transmission [13,14]. An alternative approach [15] with the potential to overcome these shortcomings involves using empirical data to characterize variability in components of the transmission cycle and applying principles of probability to combine those components to describe variability in the length of the transmission cycle as a whole—i.e., the serial interval [3].

For *P. falciparum* malaria, one analysis [7] has used such an approach to describe the generation interval, which differs from the serial interval because it pertains to the timing of infection rather than case detection. There are two important limitations of how this approach has been applied to *Pf* malaria to date, however. First, applying the generation interval to data on case data is questionable, given that the generation interval is intended to quantify the timing between infections rather than cases. Second, there are a number of heterogeneities in the *Pf* malaria transmission cycle that have not previously been incorporated into descriptions of its generation interval. We achieved a more comprehensive quantitative understanding of the *Pf* malaria generation interval and serial interval by considering 1) differences in the timing of secondary infections arising from asymptomatic or untreated cases as compared with symptomatic cases treated with antimalarial drugs, 2) variability in entomological parameters that affect the timing of transmission, 3) variability due to seasonal fluctuations in mosquito densities, and 4) inter-individual variability arising from stochastic variation in the trajectory of a given person's infectiousness over time.

## METHODS

### Overview

To obtain random variables describing the generation interval (GI) and serial interval (SI) of *Pf* malaria, we first derived random variables describing components of the GI and SI: the liver emergence period (LEP), the human-to-mosquito transmission period (HMTP), the extrinsic incubation period (EIP), the mosquito-to-human transmission period (MHTP), and the infection-to-detection period (IDP). We then summed them by direct convolution to obtain random variables describing the GI and SI. We repeated this approach under a variety of scenarios to describe variability in the GI and SI across a wide range of conditions typical of *Pf* malaria transmission in different settings.

### Probabilistic description of components of the generation and serial intervals

*Liver emergence period*. We defined the first period comprising the generation interval for *Pf* malaria as the liver emergence period. Consistent with empirical findings [16], we modeled this interval between sporozoites entering the skin and asexual merozoites emerging from the liver as a constant six days.

*Human-to-mosquito transmission period*. To simulate the trajectory of blood-stage parasites following their emergence from the liver, we used a simulation model developed by Johnston et al. [17], which tracks parasite replication beginning in the 1^st^ generation after emergence from the liver (e.g., from the 8^th^ day). Once simulated gametocytes were sufficiently mature and abundant to infect mosquitoes (sequestration time ~ Normal(7 d, 1.5 d)), we modeled the probability of a person infecting a blood-feeding mosquito as a nonlinear function of their gametocyte density, consistent with Johnston et al. [17]. Time-varying gametocytemia and its relationship with infectiousness then governed the infectiousness of a person until the infection was cleared by either the body's immune response or with the aid of antimalarial drugs. The dynamics of gametocytemia, the immune response, and the effect of antimalarial drugs were simulated with the model by Johnston et al. [18].

Because the number of mosquitoes blood-feeding on a given day can be highly variable [19], we multiplied the time-varying probabilities of infection obtained from the Johnston et al. [17] model with potentially time-varying mosquito densities, *m*(*t*). The default setting for *m*(*t*) was a constant, but for some analyses we used a time-varying function,

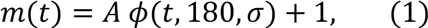
where *A* is peak amplitude, *ϕ* is a normal probability density, and σ corresponds to the width of the seasonal peak. To obtain a random variable describing the timing of a mosquito being infected by an infectious human, we multiplied the time-varying infection probabilities by *m*(*t*) and normalized the resulting curve. The sum of the LEP and HMTP for a constant *m*(*t*) is shown in Fig. 1A.

**Figure 1.**
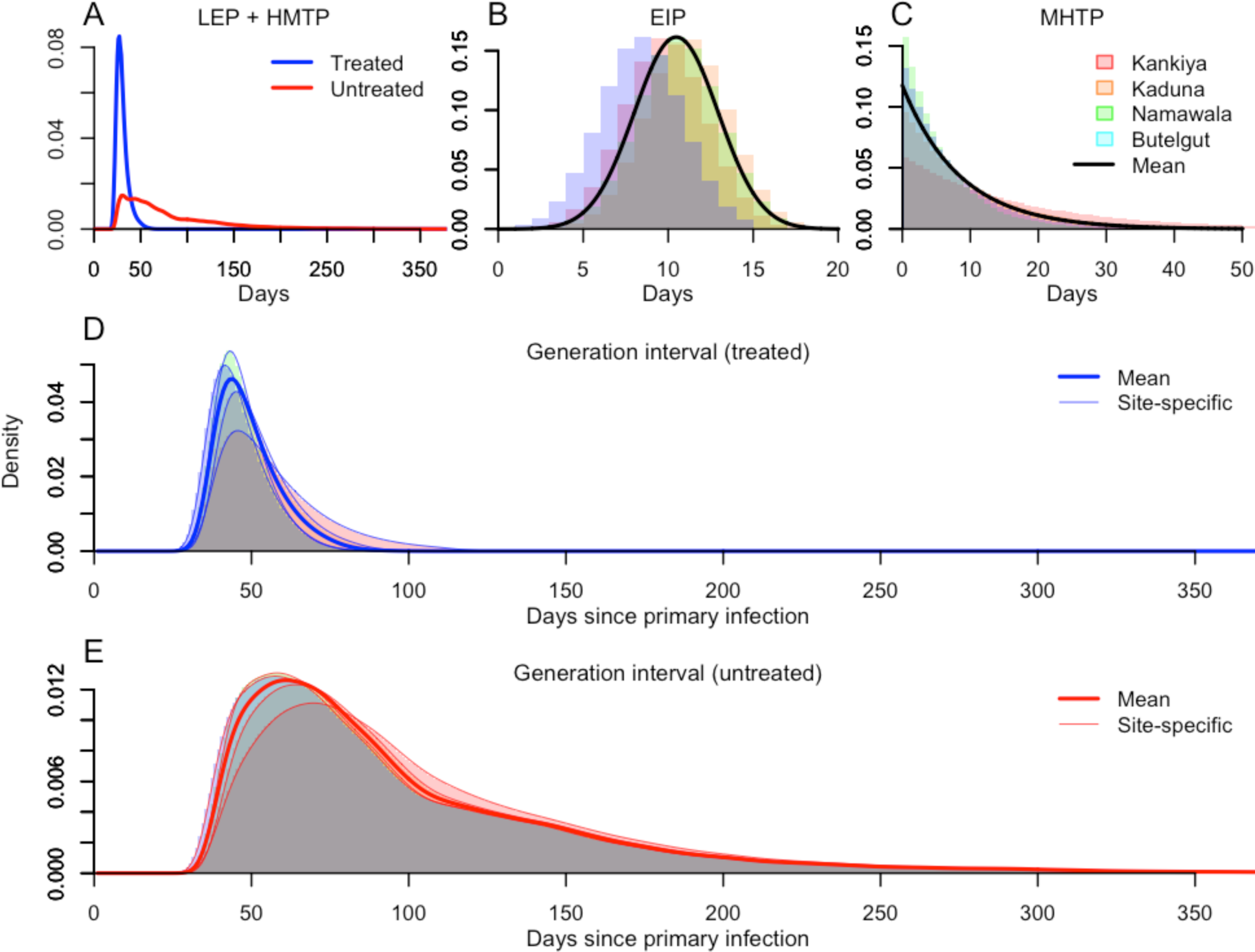
Elements of the *Pf* malaria transmission cycle (A-C) and their impact on variability in the generation interval (D, E). The first such elements that we delineated were the liver emergence period (LEP) and human-to-mosquito transmission period (HMTP), which differed for primary cases treated with antimalarial drugs or not (A). The third element was the extrinsic incubation period (EIP), whose mean values differed across four representative sites (B). The fourth element was the mosquito-to-human transmission period (MHTP), the distribution of which differed for the same four sites due to differences in mean daily mosquito mortalities (C). Combining these elements produced four site-specific generation interval distributions for treated (D) and untreated (E) primary case scenarios. Generation interval distributions with values of the entomological parameters averaged across sites are shown for comparison in D and E.

*Extrinsic incubation period*. Once *P. falciparum* gametocytes have been transmitted from an infectious human to a susceptible mosquito, a period of time known as the extrinsic incubation period (EIP) must elapse before sporozoites are produced and disseminated to the mosquito’s salivary glands, where they can then be transmitted to a human. We assumed that the EIP can be reasonably described by a normal random variable with mean estimated from any of four sites [20] and standard deviation of 2.47 days, which was estimated based on data digitized from Macdonald [21]. Because we performed our calculations on a daily basis, we modeled the EIP as a random variable with probability mass,

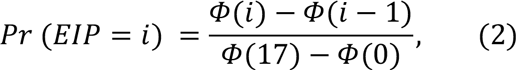
where *ϕ* is a normal cumulative density (Fig. 1B). By setting the maximum possible EIP at 17 days, eqn. 2 captures over 99% of the probability density of the corresponding normal random variable.

*Mosquito-to-human transmission period*. After a mosquito has become infectious, the final step in the transmission cycle is for the mosquito to transmit parasites to a human. To make this a tractable quantity to model, we made three simplifying assumptions. First, we assumed no senescence or any other source of variability in mortality, such that mosquitoes are subject to a constant daily probability of survival *p*. Second, we made no assumption about the feeding status of a mosquito at the time it completes the EIP and becomes infectious, and we assumed no correlation between feeding behavior and lifespan. Third, we assumed no effect of mosquito age or time since completion of the EIP on the probability of successfully infecting a human upon blood feeding. Together, these assumptions imply that the elapsed time between completion of the EIP and the time at which a human is infected can be described as a geometric random variable with probability 1 – *p*. By setting the maximum possible mosquito lifespan at 30 days beyond completion of the lowest mean EIP that we considered, we captured over 99% of the probability density of this random variable (Fig. 1C).

*Infection-to-detection period.* To calculate the serial interval distribution, we defined one additional random variable describing the interval between infection and either presentation at a clinic or detection by other means, such as active surveillance [22]. We refer to this interval as the infection-to-detection period (IDP). We modeled IDP in different ways for symptomatic (and presumably treated) and asymptomatic (and presumably untreated) cases and refer to them as IDP_S_ and IDP_A_, respectively. Common to both was the interval between sporozoites entering the skin and asexual parasites emerging from the liver, which we assumed is always six days [17].

For symptomatic cases, we added another random variable corresponding to the interval between emergence of parasites from the liver and onset of fever, which we obtained as part of the simulation output of the model by Johnston et al. [18]. The third random variable for symptomatic cases represented time elapsed before seeking treatment some number of days after the onset of fever, which we modeled as a Poisson random variable with parameter λ = 3.07. This value was obtained by maximum-likelihood estimation using data on the timing of treatment seeking relative to fever onset among 1,961 *Pf* malaria cases from Zanzibar (unpublished data).

For asymptomatic infections, we assumed that they were identified through some form of active case detection at some point during their infection when their asexual parasitemia levels exceeded 50 per μL of blood. We obtained a probability distribution describing the probability that such a level of asexual parasitemia exceeded this threshold on a given day by directly calculating the empirical density of the number of days in excess of 50 per μL from 1,000 realizations of the simulation model by Johnston et al. [17].

### Calculation of generation and serial intervals

To obtain a probabilistic description of the generation interval, we summed the LEP, HMTP, EIP, and MHTP random variables by direct convolution, resulting in

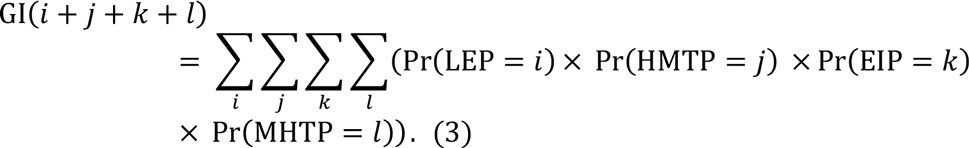

To obtain a probabilistic description of the serial interval, we summed the GI and the IDP twice, 
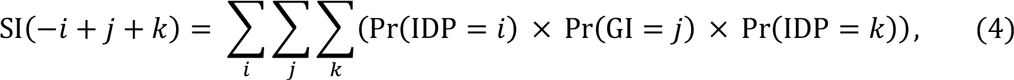
once for the primary infection and once for the secondary infection.

### Sources of variability in generation and serial intervals

Using this framework, we quantified variation in GI and SI distributions that arise from the following sources of variability in model parameters in different ecological and epidemiological contexts.

*Variability between individuals treated with antimalarial drugs or not*. To address impacts of treatment with antimalarial drugs on the generation interval of *P. falciparum* malaria, we simulated the model by Johnston et al. [17, 18] to obtain human infectivity trajectories assuming no drug treatment and assuming a standard regimen of treatment with artemisinin-based combination therapy (ACT). Treatment with ACT was modeled according to default settings in Johnston et al. [18]. In order to address variation in the lag between the manifestation of symptoms and the starting of treatment, we varied the day between the onset of fever and clinical presentation from 0-14 days, where a delay of zero days signified that the individual presented in the clinic the same day that the fever manifested. We assumed that clinical presentation marked the first day of administration of antimalarial drugs. These infectivity curves were then weighted with their respective probabilities from the Poisson distribution describing the time elapsed between fever onset and clinical presentation to arrive at a mean infectivity curve for individuals treated with antimalarial drugs.

*Geographic variability in entomological indices*. To account for variability in entomological parameters, we calculated the GI distribution under four different parameterizations of the mean EIP and daily probability of mosquito mortality corresponding to four different sites, as reported by Killeen et al. [20]. These sites and parameter values were: Kankiya, Nigeria (mean EIP = 10.3, daily mortality = 0.06); Kaduna, Nigeria (11.6, 0.10); Namawala, Tanzania (11.1, 0.17); and Butelgut, Papua New Guinea (8.9, 0.14). In cases where two estimates were reported by Killeen et al. [20], we used their average in our calculations.

*Seasonal fluctuations in mosquito densities*. To determine the extent to which seasonal fluctuations in mosquito densities could introduce variability into the GI distribution, we set *m*(*t*) equal to the time-varying function in eqn. 1 and calculated the resulting GI and SI distributions. We performed these calculations under four scenarios about the parameters in eqn. 1: *A* = 1, σ = 14; *A* = 9, σ = 14; *A* = 1, σ = 120; and *A* = 9, σ = 120. Values of *A* equal to 1 and 9 led to two- or ten-fold increases, respectively, in the ratio of high-season to low-season mosquito densities. Values of *σ* of 14 or 120 lead to narrow or wide seasonal peaks in mosquito density, respectively.

*Inter-individual variability in gametocytemia trajectories*. To assess the extent of possible variability in different GI and SI distributions among different individuals with different realized HMTP distributions, we simulated 1,000 realizations of HMTP distributions from the model by Johnston et al. [17, 18]. We compared these against our default HMTP distribution, which we obtained by taking the mean of these realizations.

## RESULTS

We first analyzed differences in generation interval (GI) distributions arising from treated and untreated primary cases (Fig. 1A). The overall shape of the GI distribution was dependent on the status of the primary case with respect to antimalarial drug treatment, with mean (SD) of 49.1 (10.2) and 101.6 (62.2) days for GIs arising from treated and untreated primary cases, respectively (Fig. 1D,E). Overall, GIs arising from untreated primary cases were much longer, with 95% of secondary cases being infected by day 68 for treated primary cases as opposed to 219 days for untreated primary cases.

We then calculated serial interval (SI) distributions and compared four different versions with different combinations of treated and untreated primary cases and symptomatic and asymptomatic secondary cases. The first step in this process involved calculating probabilistic estimates of the infection-to-detection period for treated (symptomatic) and untreated (asymptomatic) primary (secondary) cases. The former, IDP_S_, was relatively short (16.6 (3.1), Fig. 2A), whereas the latter, IDP_A_, was relatively long and leptokurtic (69.8 (48.8), Fig. 2B). Relative to the timing of primary cases presenting at a clinic and treated with drugs, symptomatic secondary cases would be expected to appear 49.1 (11.1) days later (Fig. 2C) and asymptomatic secondary infections detected through active case detection would be expected to appear 102.2 (49.9) days later (Fig. 2D). Relative to the timing of asymptomatic primary infections, secondary cases presenting clinically would be expected to appear 48.4 (79.1) days later (Fig. 2E) and asymptomatic secondary infections detected by active case detection would be expected to appear 101.6 (92.9) days later (Fig. 2F). These distributions displayed some sensitivity to the choice of the asexual parasitemia threshold for detection (Fig. 3), but this sensitivity was small for thresholds within an order of magnitude range (10–100 asexual parasites per μL of blood). Additionally, we note that 24.5% of secondary cases presenting clinically are expected to do so prior to the associated primary infection being detected by active case detection (Fig. 2E), assuming that the primary infection is ever detected. Some 11.5% of asymptomatic secondary infections detected by active case detection could be detected prior to detection of the associated primary infection (Fig. 2F).

**Figure 2.**
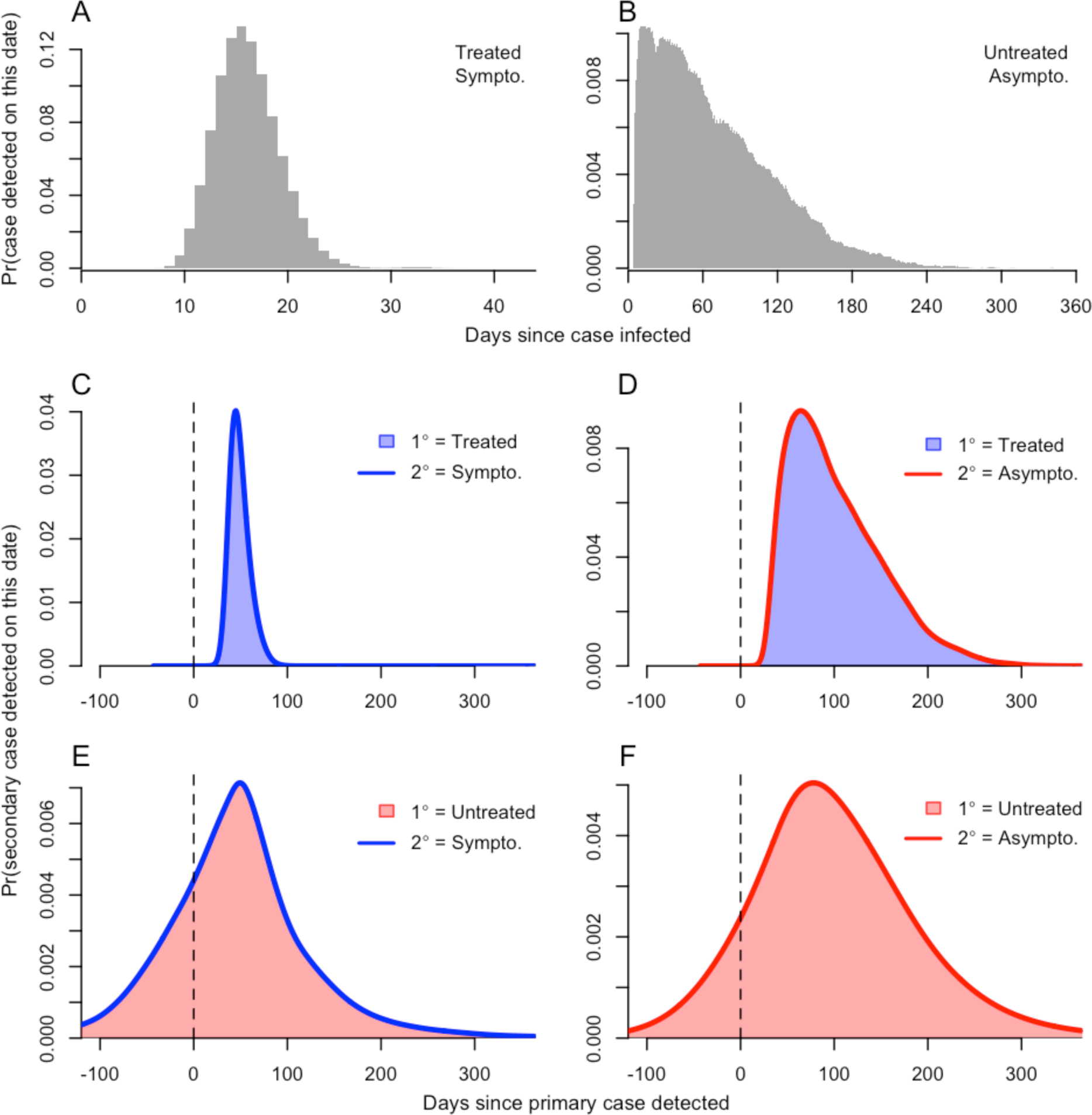
Probabilistic descriptions of the infection-to-detection periods IDP_S_ and IDP_A_ between infection and detection of treated / symptomatic (A) and untreated / asymptomatic (B) cases, respectively. Combining IDP random variables with the appropriate generation interval random variables yielded four different estimates of the serial interval: treated primary case and either symptomatic (C) or asymptomatic (D) secondary case; untreated primary case and either symptomatic (E) or asymptomatic secondary case (F).

**Figure 3.**
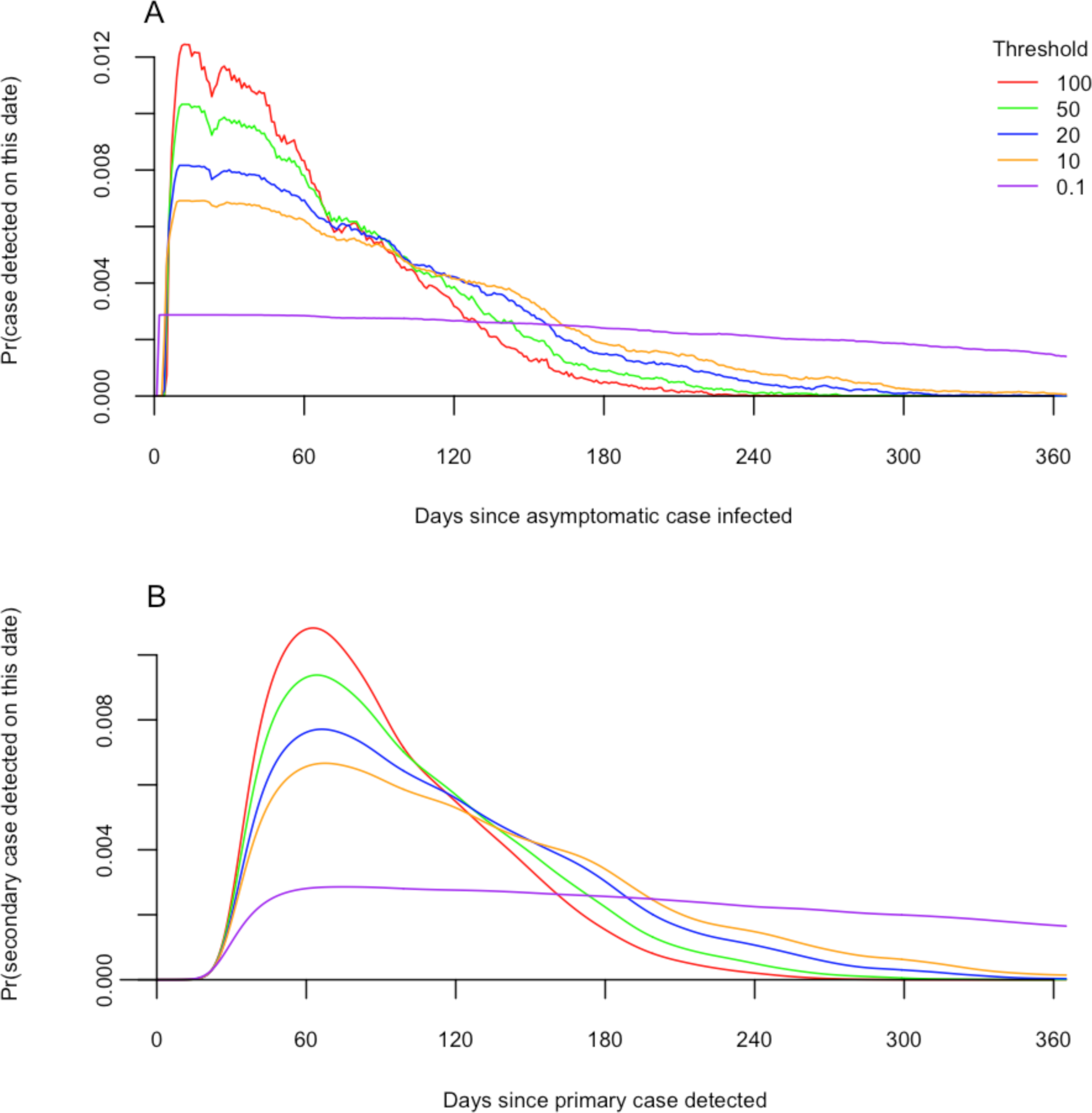
Variability associated with differences in the asexual parasitemia detection threshold for a secondary infection detected by active case detection. Panel A shows normalized probability densities for the infection-to-detection period IDP_A_ as a function of the detection threshold (asexual parasites per μL of blood). Panel B shows normalized probability densities for the serial interval given an untreated primary case as a function of the corresponding IDP_A_ distributions from A.

Using entomological parameters from four diverse sites (Fig. 1B,C), means and standard deviations of the GI distributions varied from 46.1 (9.3) to 56.6 (16.2) for treated primary cases (Fig. 1D) and from 98.7 (62.1) to 109.2 (63.5) for untreated cases in Butelgut and Kankiya, respectively (Fig. 1E). By comparison, the modes of the GI distributions ranged 42–46 for treated primary cases and 58–70 for untreated cases (Fig. 1D & 1E). The long GI for Kankiya appears to be driven mostly by very low mosquito mortality, and the short GI at Butelgut appears to be driven by both a short EIP and relatively high mortality (Fig. 1B,C).

Differences in GI distributions owing to differences in timing relative to a seasonal transmission peak were more substantial, affecting not only the moments of the GI distribution but also its shape (Fig. 4). These effects were most pronounced for untreated primary cases, whose GI distributions spanned a broader portion of the year (Fig. 4, right column). Primary cases infected well before the seasonal peak tended to be associated with more secondary cases later than they would have in a constant environment, and the peak of the GI distribution for primary cases infected just before the seasonal peak tended to be more peaked and narrower than it would have otherwise. The extent of these differences depended on the extent to which transmission was seasonally peaked (Fig. 4, second row).

**Figure 4.**
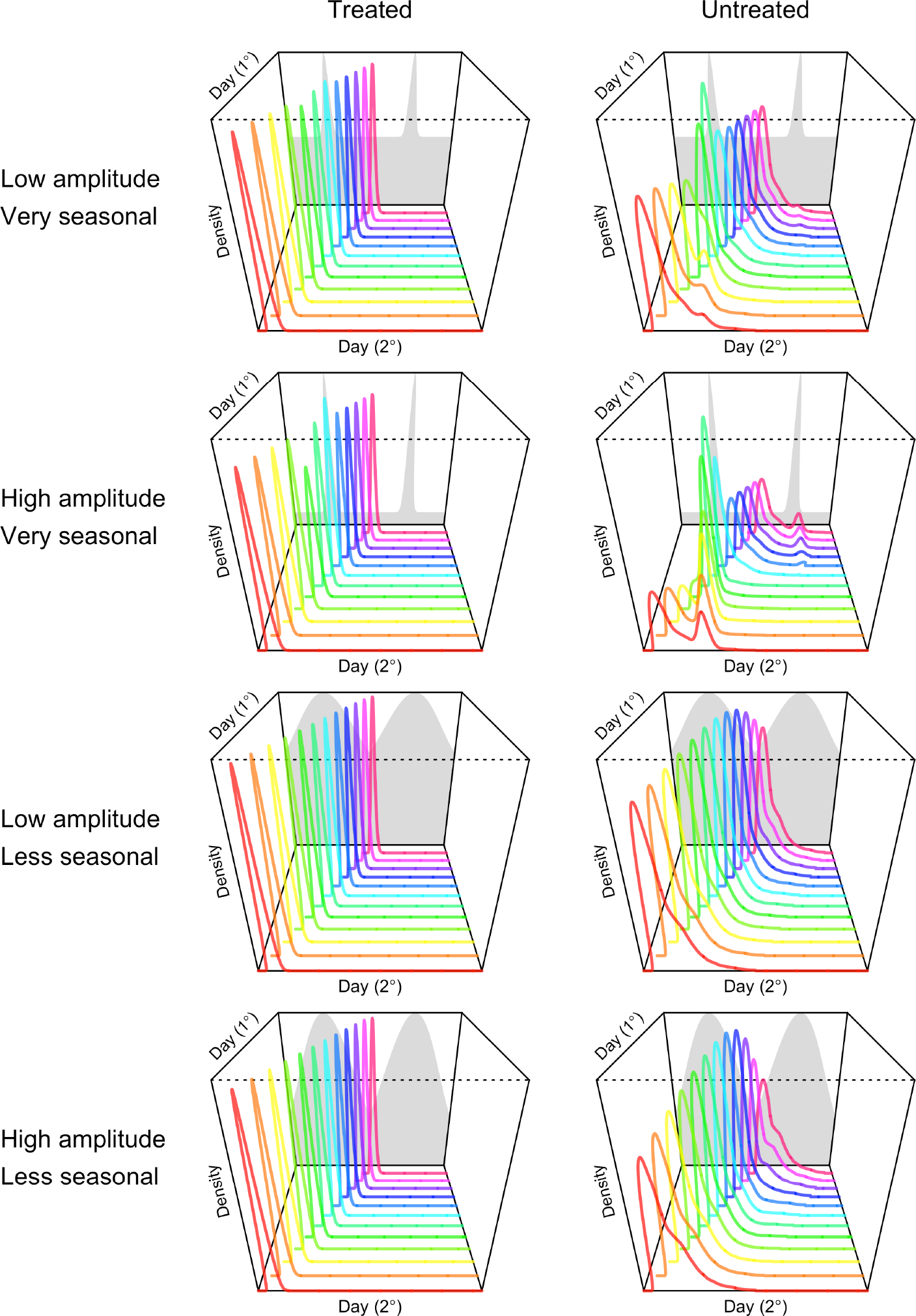
Variability in the generation interval distribution in a seasonal environment for treated and untreated primary cases (columns) in seasonal environments with different properties (rows). Seasonality was imposed by forcing mosquito densities consistent with the gray shapes in the background of each panel, which vary in their amplitude and the distinctiveness of the seasonal peak at day 180 in each of two years. Serial interval distributions are shown for primary infections occurring on days 1 through 360 in increments of 30.

The final source of variability in the GI distributions that we examined pertained to variability in the timing of infectiousness of humans to mosquitoes across 1,000 simulated primary cases with the same drug treatment status. For primary cases receiving antimalarial drugs, the probabilistic descriptions of the human-to-mosquito transmission period (HMTP) across different individuals were relatively uniform, with all individual trajectories that we simulated rising and falling relatively quickly (Fig. 5 A,B). For primary cases not receiving antimalarial drugs, probabilistic descriptions of the HMTP across different individuals were much more variable. Unlike treated cases, untreated cases displayed simulated HMTP distributions with multiple peaks; the number, timing, and height of which varied considerably (Fig. 5C). These differences lead to broad variability in quantiles of the GI distribution. For example, the median GI varied by over 100 days for the inner 95% of individual GI distributions.

**Figure 5.**
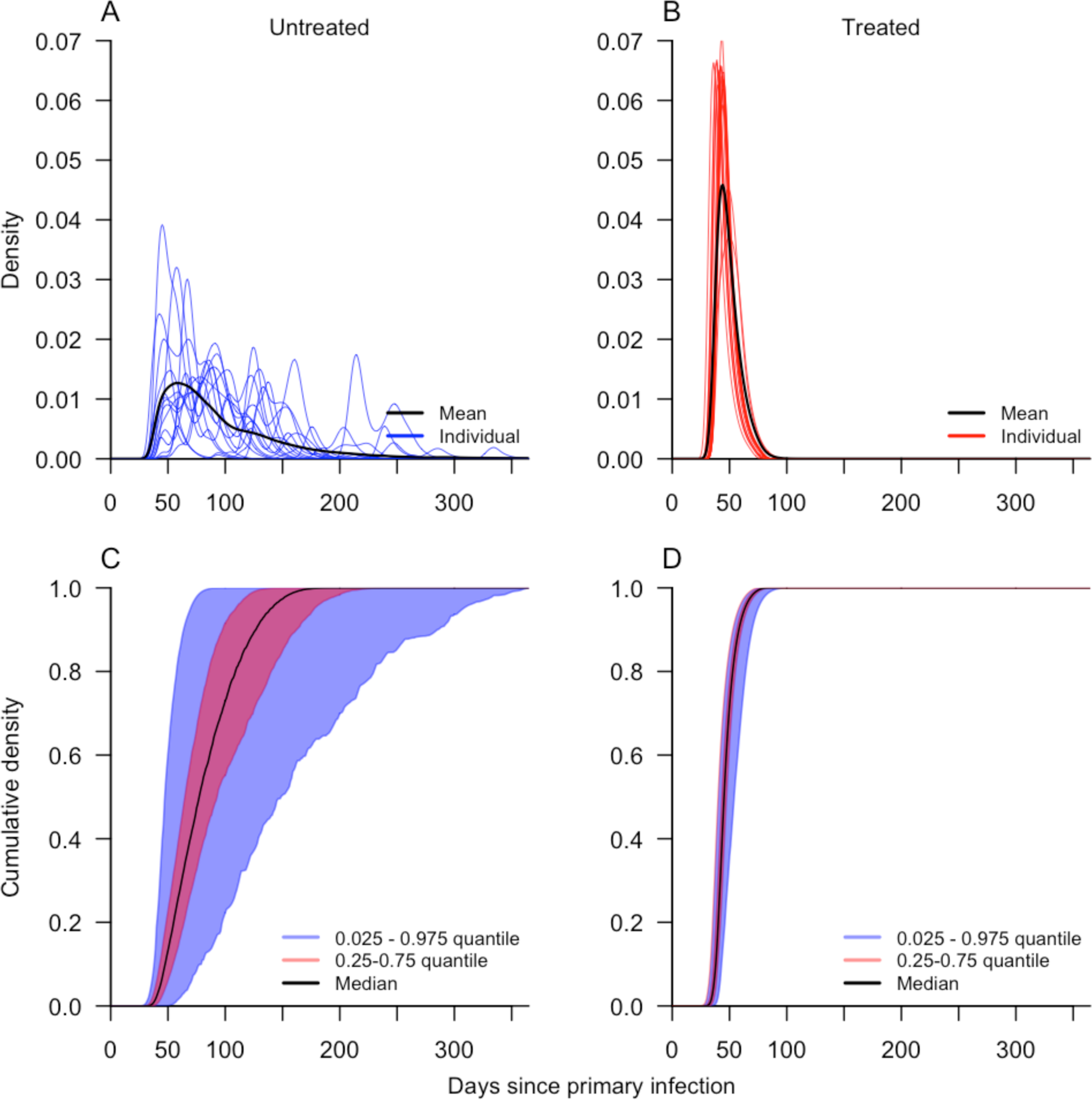
Variability in generation interval distributions ensuing from primary infections that do (right) or do not (left) receive antimalarial drugs. The top row shows normalized probability densities for mean and representative generation interval distributions from 15 realizations of the simulation model. The bottom row shows quantiles of cumulative probability densities for 1,000 realizations of the simulation model.

## DISCUSSION

One of the first attempts to quantify the serial interval for *Pf* malaria was by Macdonald [23], who posited that 36 days represents a minimum estimate based on hard biological constraints. Much more recently, Churcher et al. [7] posited a mean generation interval of 102 days for individuals who do not receive antimalarial drug treatment and 33 days for those who do. Our estimates are in good agreement with the former (102 days) but not the latter (48 days). One reason for this nearly 50% discrepancy has to do with differences in our assumptions about the delay between onset of symptoms and seeking of treatment. Although their means were nearly identical, the exponential distribution used by Churcher et al. [7] resulted in appreciably more individuals seeking treatment on the same day as symptom onset than our Poisson distribution did. Collectively, this and other seemingly subtle differences led to a large discrepancy between the means of our and Churcher et al.'s [7] distributions for treated infections. The discrepancy between the distributions for untreated infections was smaller due to the dominance of a much lengthier period of human infectiousness.

An increasingly important application of probabilistic descriptions of the serial interval is the inference of transmission linkages between cases [5–7, 24]. Given the breadth of the serial interval distributions that we calculated, we conclude that using temporal information alone to link *Pf* malaria cases may be inadvisable. First, even in the best-case scenario of a putative transmission linkage between two known cases that promptly sought treatment, the serial interval distribution is sufficiently wide that distinguishing that linkage from others within a time period of a few weeks should be largely uninformed by temporal data alone. Second, the shape of the generation interval distribution differs considerably from person to person due to the complex within-host dynamics of *Pf* infections [6, 25]. This may preclude the inference of transmission linkages ensuing from primary cases whose infections do not follow average behavior. Third, serial interval distributions associated with cases that were or were not treated with antimalarial drugs differ substantially, with the latter even being negative in many cases (i.e., the secondary case is detected before the primary case is detected). Given that negative values are strictly impossible for generation intervals, this underscores the importance of being conscientious about the distinction between generation and serial intervals when applying these methods to case data (as in [6]).

Probabilistic descriptions of generation and serial intervals also have an important role to play in population-level models of infectious disease dynamics. Together with an estimate of epidemic growth rate, the generation interval distribution can be used to estimate the basic reproduction number and related quantities [8, 26, 27]. Any time that there are secular changes in factors that affect transmission within the timeframe of a single generation, however, there is a risk of being misled by a static description of the generation or serial interval distribution.Similar to our analysis of how seasonally varying mosquito densities affect the *Pf* malaria generation interval, an analysis by Vynnycky and Fine [28] showed that not accounting for secular trends in contact rates over time lead to an underestimate of tuberculosis transmission potential. This result underscores our conclusion that there is no one-size-fits-all description of generation and serial intervals, particularly for long-lasting infections such as *Pf* malaria.

## CONCLUSIONS

We have highlighted a number of reasons why generation and serial interval distributions are variable for *Pf* malaria and have offered quantitative remedies to many of those situations. To this end, code for reproducing our figures and for calculating generation and serial interval distributions over one or more generations of *Pf* malaria cases is available at https://github.com/TAlexPerkins/malaria_serial_interval. Like many topics in epidemiology, robust quantification of generation and serial interval distributions stands to benefit from careful and empirically well-grounded use of mechanistic models to describe constituent processes in the transmission cycle.

## ACKNOWLEDGEMENTS

This work was supported by the Research and Policy for Infectious Disease Dynamics (RAPIDD) program of the Science and Technology Directory, Department of Homeland Security, and Fogarty International Center, National Institutes of Health. BG and DLS received support from a grant from the National Institutes of Health (ICMER U19 AI089674). BG and TAP received support from a grant from the Bill and Melinda Gates Foundation (OPP 1132226 to BG), as did DLS and TAP (OPP 1110495 to DLS).

